# Amelioration of Oxidative Stress in Sodium Fluoride-administered Wistar Rats by *Tamarindus indica* Fruit Pulp Ethanolic Extract

**DOI:** 10.1101/2024.10.01.615996

**Authors:** Lutfat A Usman, Emmanuel O. Ajani, Muhammed A Ishiaku, Taofeeq A Bankole, Rasheed B. Ibrahim, Dauda K. Saka, Hassan T. Abdulameed, Raliat A. Aladodo

## Abstract

Infertility is a significant global health issue often exacerbated by environmental toxins like sodium fluoride (NaF), which induces oxidative stress and reproductive dysfunction. Medicinal plants, such as Tamarindus indica, known for their antioxidant properties, offer potential natural remedies. This study evaluates the antioxidant potential of the ethanolic extract of Tamarindus indica fruit pulp in mitigating NaF-induced infertility in Wistar rats.

*Tamarindus indica* fruits were authenticated and processed into a crude extract, with phytochemical screening revealing the presence of tannins and steroids. Seventy Wistar rats were divided into seven groups, receiving various treatments including T. indica extract, NaF, and Clomid. The acute toxicity test, conducted according to WHO and OECD guidelines, showed no lethal effects up to 5000 mg/kg of the extract. Phytochemical screening indicated that the antioxidant properties of the extract could be attributed to tannins and steroids.

Experimental groups treated with T. indica extract exhibited significant improvements in oxidative stress markers such as malondialdehyde (MDA), glutathione peroxidase (GPx), glutathione (GSH), catalase (CAT), and superoxide dismutase (SOD) compared to control groups. The 200 mg/kg preventive group demonstrated the most notable antioxidant activity, comparable to the normal control group.

The findings suggest that the ethanolic extract of Tamarindus indica fruit pulp at 200 mg/kg body weight has significant potential as a natural therapeutic agent against oxidative stress-induced infertility. This highlights its promise as an alternative to conventional treatments, offering a safe and effective option for managing reproductive health issues related to oxidative stress.

## INTRODUCTION

Infertility is defined as the inability to conceive naturally after one year of regular unprotected intercourse (Taylor, 2003). For healthy young couples, the likelihood of getting pregnant varies. An estimated 48.5 million couples worldwide were infertile in 2010 (Mascarenhas *et al*., 2012). Most of the infertile couples have one of three major causes including a male factor, ovulatory dysfunction, or tubal-peritoneal disease.

In humans, oxidative stress is thought to be involved in the development of cancer (Halliwell, 2007), parkinson’s disease (Hwang, 2013), Lafora disease (Roma-Mateo *et al*., 2015), alzheimer’s disease, atherosclerosis, heart failure, myocardial infarction, fragile X syndrome, sickle-cell disease, lichen planus, vitiligo, autism, infection, chronic fatigue syndrome, and depression (Kennedy *et al*., 2005; Jimenez-Fernandez, 2015). However, reactive oxygen species can be beneficial, as they are used by the immune system as a way to attack and kill pathogens. Short-term oxidative stress may also be important in prevention of aging by induction of a process named mitohormesis (Jimenez-Fernandez, 2015). Of particular interest to this study is the report that sperm DNA fragmentation appears to be an important factor in the aetiology of male infertility. Thus, men with high DNA fragmentation levels have significantly lower odds of conceiving. Oxidative stress is the major cause of DNA fragmentation in spermatozoa (Wright et al., 2014). A high level of the oxidative DNA damage 8-OHdG is associated with abnormal spermatozoa and male infertility (Guz *et al*., 2013).

Tamarind (*Tamarindus indica*) is a leguminous tree in the family Fabaceae that is indigenous to tropical Africa. It is botanically identified as a *Tamarindus indica Linn*., the member of Caesalpinaceae subfamily of Fabaceae family. The tree averages 20-25 m in height and 1 m in diameter, slow growing, but long lived, with an average life span of 80-200 years (El-Siddig *et al*., 1999). The tamarind tree produces edible, pod-like fruits which are used extensively in cuisines around the world. Virtually every part of *Tamarindus indica* (wood, root, leaves, bark and fruits) has either nutritional or medicinal value, with a number of industrial and commercial applications (Emmy *et al*., 2010). Tamarind is useful traditionally in gastric disorders, bilious vomiting, scurvy, datura poisoning, alcoholic intoxication, scabies, pharyngitis, otalgia, stomatitis, constipation, haemorrhoids and eye diseases. Tamarind pulp is also said to aid in the cure of malarial fever (Timyan, 1996). The fruits are reported to have hypo-lipidemic, anti-inflammatory, anti-fungal and anti-bacterial properties (John *et al*., 2004). Tamarind pulp typically contains 20.6% water, 3.1% protein, 0.4% fat, 70.8% carbohydrates, 3.0%fibre and 2.1% ash (El Siddig *et al*., 1999). Tamarind seeds are traditionally used to treat diabetes, fevers and intestinal infections. The stem and bark also have medicinal properties (Naveena *et al*., 2012). Tamarind has been used for centuries as a medicinal plant; its fruits are the most valuable part which has often been reported as curative in several pharmacopoeias.

This study aimed to investigate the ameliorative effect of *Tamarindus indica* fruit pulp extract on oxidative stress markers in sodium fluoride-induced reproductive dysfunction in Wistar rats.

## MATERIALS AND METHODS

### Drug Sample and chemicals

Clomifene citrate (clomid) was a product of *Bruno Farmaceutici* S.p.A and was purchased at Farason Pharmaceuticals Ilorin, Kwara State. Other chemicals and kits used in this study were obtained from Merck (India) and Sigma-Aldrich (Saint Louis, Missouri).

### Plant material

#### Collection and Authentication

The compact fruit of *Tamarindus indica* were obtained from farm land around Kwara State University, Malete campus in the month of September, 2018. The plant material was identified and authenticated by a botanist at the Herbarium of the Department of Plant Biology, University of Ilorin, Nigeria, where a voucher specimen with number UILH/001/1155 for *Tamarindus indica* was deposited.

#### Extraction

Dust free clean pulps were dried in an oven at 45^0^C. The dried fruit (916.77g) was then macerated in 2338 ml of 70% ethanol for 72 hours. The resulting macerate was blended to smoothness using a high-speed electric blender. Afterwards, the blended macerate was filtered using a muslin cloth, cotton-wool plug and Whatman No. 1 filter paper respectively. The resulting filtrate was concentrated to dryness using a vacuum oven at 60°C until a constant weight was achieved. The crude extract was stored in an air tight container and was placed in the refrigerator at a temperature of 4ºC prior to use.

### Experimental Protocols

#### Experimental Animals

Seventy clinically healthy rats of both sexes (in same proportion) with an average weight of 90 ± 5 g were obtained from Umaraahmarkeen Nigeria Global Ventures, Kwara State, Nigeria. The animals were maintained in stainless steel cages at the animal holding unit of the Department of Medical Biochemistry and Pharmacology, Kwara State University, Malete with optimum environmental conditions (temperature: 23 ±1°C, photoperiod: 12 h light/dark cycle, relative humidity: 45-50%). The animals were allowed free access to food and fresh water *ad libitum*. All the animals were acclimatized to laboratory conditions for a week before commencement of the experiment. The study was executed following approval from the Ethical Committee on the use of Laboratory Animals of Department of Medical Biochemistry and Pharmacology, Kwara State University, Malete, Nigeria and management and maintenance was in accordance with the principles of laboratory animal care (NIH publication no. 82-23, revised 1996) guidelines.

#### Induction of Infertility

Experimental rats were rendered infertile after 45 days of continuous oral administration of 20 mg/kg of sodium fluoride (NaF) daily (Zhou *et al*., 2013; Zhang *et al*., 2016).

#### Animal grouping and Treatments

Seventy (70) rats were randomized into seven groups of 10 rats (5 males and 5 females) per group. The rats were labelled and treated as follow: Control (NC): non induced and administered with normal saline; Model group(MG): administered with NaF and normal saline; Preventive Group 1(PG1): first administered with 200 mg/kg b.w of *Tamarindus indica* fruit extract for 4 weeks followed by concurrent administration of NaF for another 45 days; Preventive Group 2 (PG2): first administered with 400 mg/kg b.w of *Tamarind indica* fruit extract for 4 weeks followed by the concurrent administration of NaF for the next 45 days; Treatment Group 1 (TG1): initially administered with NaF for the first 45 days after which they were concurrently treated with 200 mg/kg b.w of *Tamarindus indica* fruit extract for another 4 weeks; Treatment Group 2 (TG2): initially administered with NaF for 45 days and thereafter simultaneously treated with 400 mg/kg b.w of *Tamarindus indica* fruit extract for the next 4 weeks; Positive Control (PC): first administered with NaF for 45 days and then simultaneously treated with Clomid for another 4 weeks. NaF was administered at 20 mg/kg and clomid at 50mg/kg. All treatments were administered orally as a single dose per day using an oral intubator every day. Twenty-four hours after the last treatment, under mild diethylether anesthesia, the animals were sacrificed and blood was obtained from the jugular vein. Blood sample was transferred into plain centrifuge tube and allowed to clot at room temperature. It was then centrifuged within 1 hour of collection at 4000x g for 10 minutes to obtain the serum.

### Biochemical assay

#### Determination of GSH, CAT, GPx, MDA and SOD

The assay was carried out according to the method described by Sun *et al*., (2007). The assay employs the quantitative sandwich enzyme immunoassay technique. Antibody specific for GSH-Px/CAT/SOD/MDA/GSH was pre-coated onto a microplate. Standards and samples were pipetted into the wells and any GSH-Px/CAT/SOD/MDA/GSH present was bound by the immobilized antibody. After removing any unbound substances, a biotin-conjugated antibody specific for the biomarker was added to the wells. After washing, avidin conjugated Horseradish Peroxidase (HRP) was added to the wells. Following a wash to remove any unbound avidin-enzyme reagent, the substrate solution was added to the wells and colour develops in proportion to the amount of protein bound in the initial step. The colour development is stopped and the intensity of the colour is measured.

#### Determination of malondialdehyde

The assay was carried out according to the method described by Sun *et al*., (2007). The assay employs the colorimetric technique. In the lipid peroxidation assay protocol, the MDA in the sample reacts with thiobarbituric acid (TBA) to generate a MDA-TBA adduct. TBA solution was added to samples and standards and incubate at 95ºC for 60 min. The solution was cooled in ice bath for 10 min and afterwards, transferred to wells of microplate and the MDA-TBA adduct was quantified colorimetrically (OD = 532 nm).

#### Statistical Analysis

All data were expressed as the mean of five replicates + standard error of mean (S.E.M). Statistical evaluation of data was performed using SPSS version 16.0 using one way analysis of variance (ANOVA), followed by Duncan’s posthoc test for multiple comparison. Values were considered statistically significant at *p* < 0.05 (confidence level = 95%).

## RESULTS

The malondialdehyde level of the male control rats was significantly (p<0.05) lower when compared with the female counterparts. However, when the GSH level of the model groups were compared, the model male rats showed an elevated GSH level above that of the female group. Administration of NaF in the male rats resulted in significant increase in the malondialdehyde above the control value (Figure 1). The malondialdehyde value obtained for the male rats treated with clomid was not significantly different from that of the model group. All the *Tamarindus indica* treated male rats showed malondialdehyde level that were significantly lower than that of the model group. The value obtained for the rats pre-treated with 200 mg/kg dose of the extract was not significantly different from that of the control value. The results of the malondialdehyde value obtained for the female rats showed that administration of NaF significantly reduced malondialgehyde level in the model group. The result also showed that when pre-treated with the extract and following treatment after NaF administration, the level of malondialdehyde increased above the model group value to a value not significantly different from that of the control value.

**FIGURE 1.**
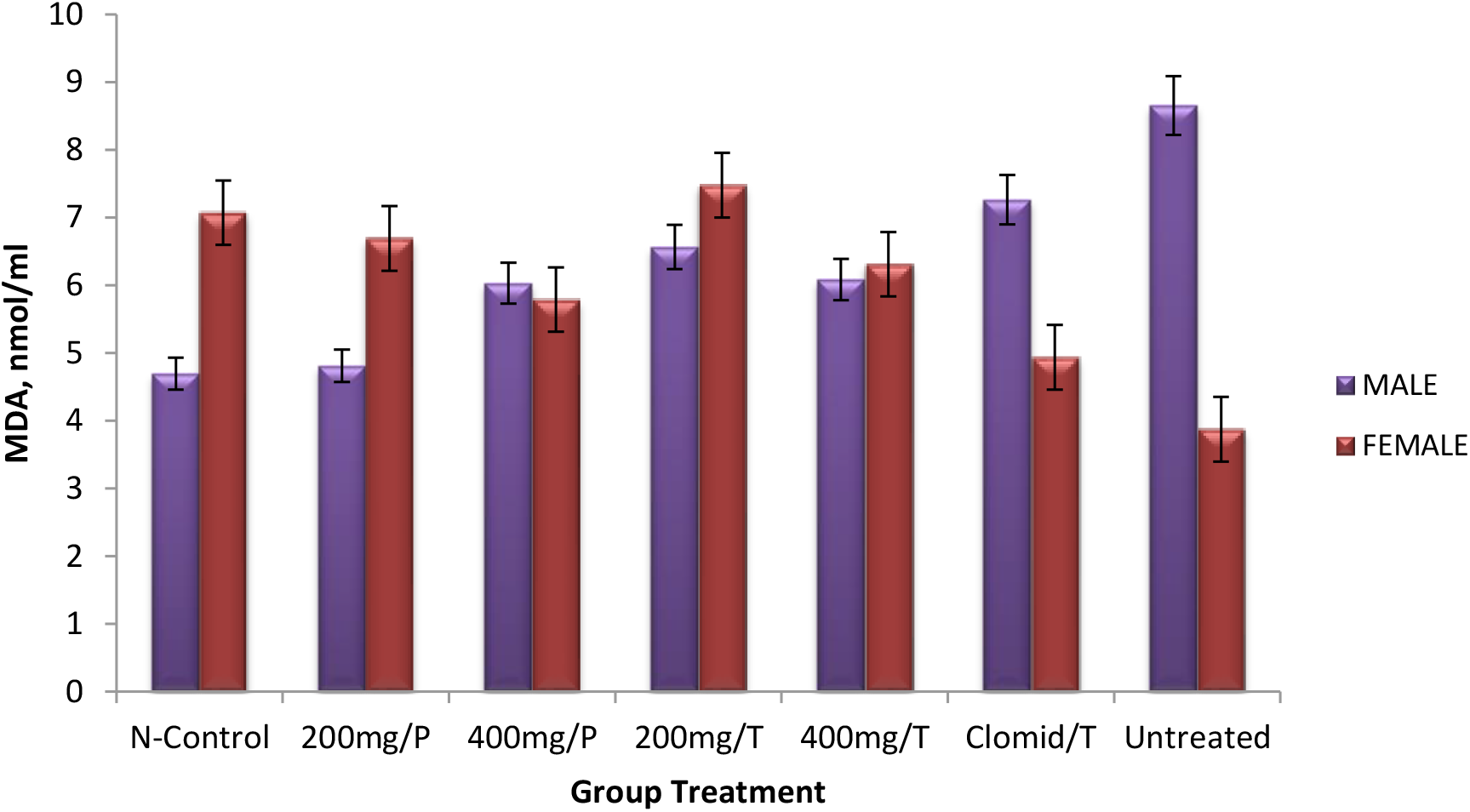
Effect of *Tamarindus indica* fruit pulp extract on serum malondialdehyde (MDA) levels in Wistar rats. Results are Mean ±S.E.M, (n=5).

The GSH level of the male control rats was observed to be significantly higher when compared with that of the female counterparts (Figure 2). The GSH level of the model male rats was also observed to be higher than that of the female group. When compared among the treated groups, the GSH level of all the treated male rats were not significantly different from each other. They were also not significantly different from the control value. Similarly, no significant difference was observed among the GSH level of the extract treated female rats when compared among each other. They were also not observed to be different from that of the control group.

**FIGURE 2.**
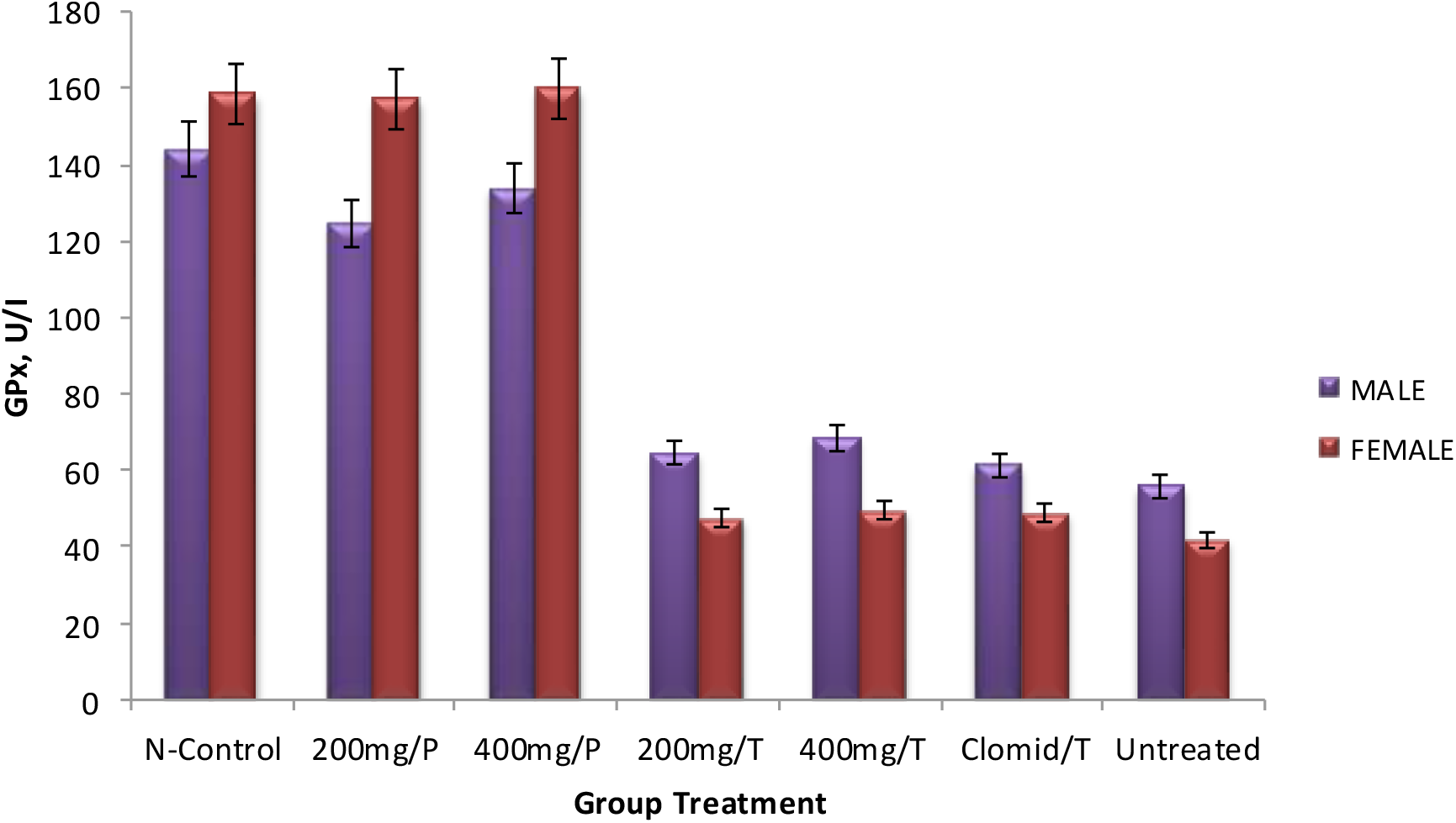
Effect of *Tamarindus indica* fruit pulp extract on serum glutathione peroxidase (GPx) levels in Wistar rats. Results are Mean ±S.E.M, (n=5). Bars with different letters are significantly different (*p* 0.05).

The result of the serum glutathione peroxidase showed that following administration of NaF, in both the male and female rats, there were significant reductions in the glutathione peroxidase activities (Figure 3). The Figure also showed that when treated with *Tamarindus indica* extract (at both doses) and with clomid, the glutathione peroxidase activities (in all the rats) were not significantly different from that of the model groups but were lowered than that of the control group. Pre-treatment with the extract however, showed a significant increase in the gutathione peroxidase activities above that of the model group. The observed glutathione peroxidase activity in the rats pre-treated with 200 mg/kg dose of *Tamarindus indica* extract was above that of the control value.

**FIGURE 3.**
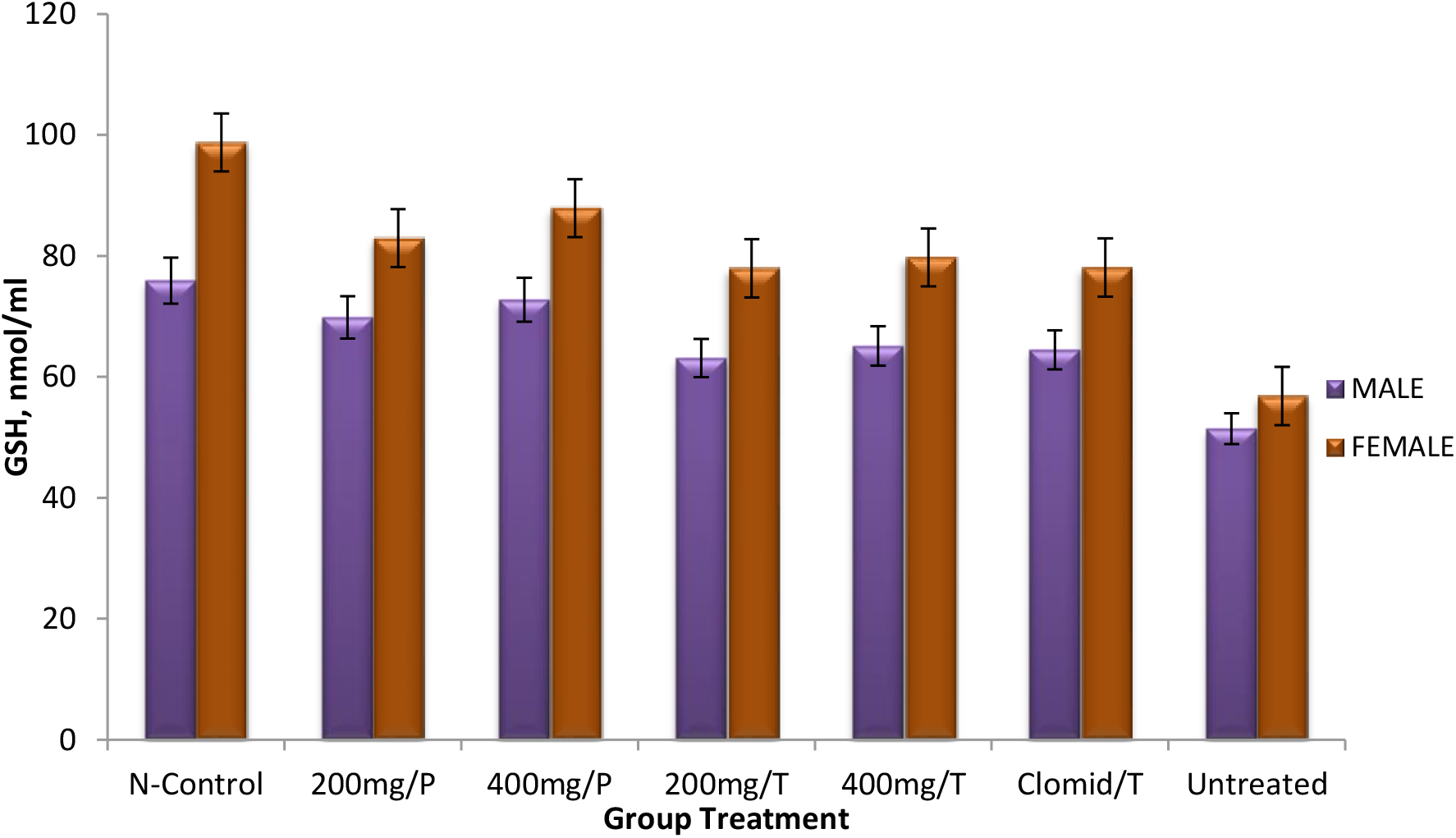
Effect of *Tamarindus indica* fruit pulp extract on serum glutathione (GSH) levels in male Wistar rats. Results are Mean ±S.E.M, (n=5).

**FIGURE 4.**
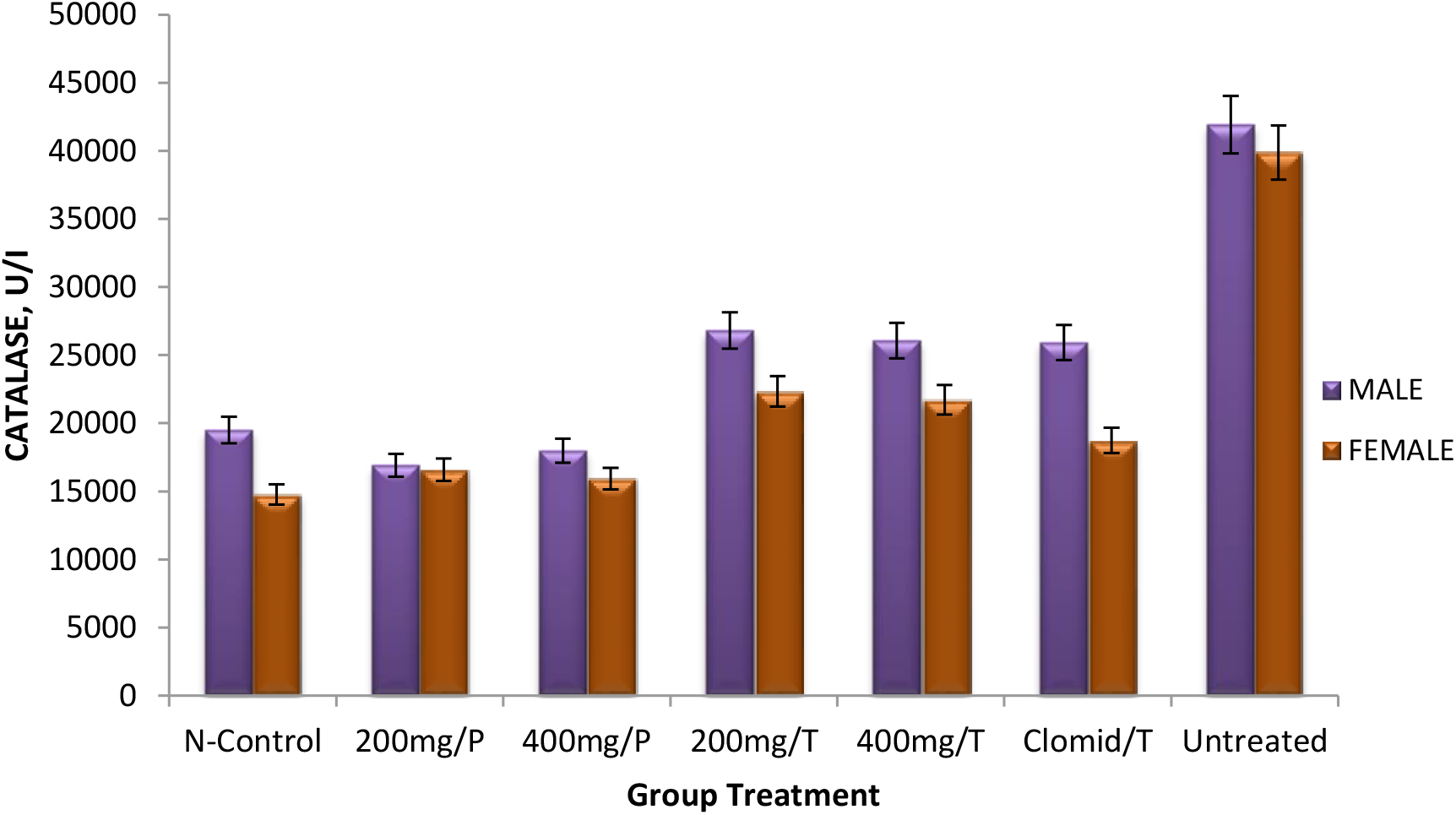
Effect of *Tamarindus indica* fruit pulp extract on serum catalase (CAT) levels in Wistar rats. Results are Mean ±S.E.M, (n=5).

Serum catalase activities of the male control rats was not significantly different from that of the female group (Figure 5). Similarly, no significant different was observed between the catalase activities of the model male rats and that of the female counterpart. Pre-treatment with *Tamarindus indica* extract at both doses showed catalase activities in the male rats that were significantly lowered than that of the model group. Although the observed activity in the 200 mg/kg treated group was not different from that of the control group, the observed activity in the 400 mg/kg treated group was higher than that of the control group. The observed catalase activities in the male rats treated *Tamarindus indica* after the NaF administration were significantly lower than that of the model group and that of the control group. The female rats pre-treated with the extract showed catalase activities that although lowered than that of the model group, but were however not different from the control value. Treatment with the extract after NaF administration showed catalase activities that were also lower than that of the model group and were also lower than that of the control group.

**FIGURE 5.**
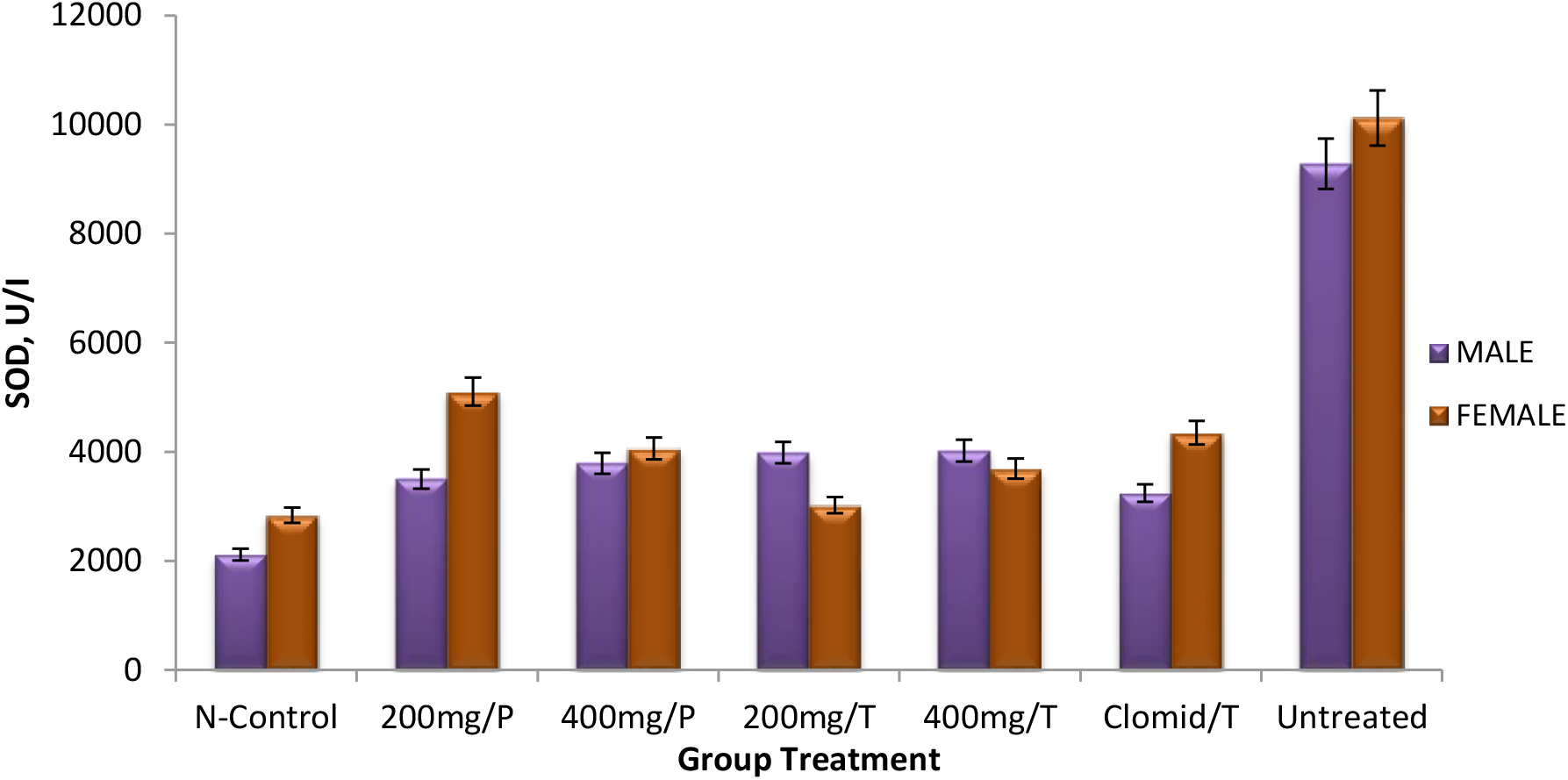
Effect of *Tamarindus indica* fruit pulp extract on serum superoxide dismutase (SOD) in Wistar rats. Results are Mean ±S.E.M, (n=5).

The result of the serum superoxide dismutase activities (Figure 5) showed that the superoxide dismutase activities of the male control rats were not different from that of the female group. Similarly, no significant difference was seen between the activities of the enzyme in the model male rat and the female counterpart. Pre-treatment of the male rats with the extract at both doses and treatment after NaF administration raised the superoxide dismutase activities above that of the control group although the observed activities were significantly lower than that of the model group. Similarly, pre-treatment of the female rat with the extract and treatment after NaF administration also raised the superoxide dismutase activities above that of the control value although the observed activities were observed to be also lower than that of the model group.

## DISCUSSION

Malondialdehyde, a product of lipid peroxidation, is often used as a marker of oxidative stress (Nair *et al*., 2008). We observed in the study that there was an increased malodialdehyde level in male rats compared to the female group indicating high level of lipid peroxidation in male rats than the female rats. This is in line with the findings of Yamaguti *et al*. (2013) and Morales-Gonzalez *et al*. (2010). The result of the present study showed am elevation in malondialdehyde level following induction of reproductive dysfunction in male rats whereas a reduction in the same parameters were observed in the female reproductive dysfunctioned group. The result suggests that whereas reproductive dysfunction in male rats may be associated with increased lipid peroxidation, a decrease lipid peroxidation may be associated with reproductive dysfunction in the female. In an earlier study, Yamaguti *et al*. (2013) reported an increase in serum malondialdehyde levels in rats (both sexes) administered continuously with NaF at varying concentrations. Our observation that administration of the extract lowered the MDA level in the male rats below that of the model group indicates that *T. indica* is able to effectively reduced lipid peroxidation associated with male reproductive dysfunction. This is in agreement with the findings of Spittle (2008). This was however different from observation with female rats where treatment with the extract was observed to increase the level of lipid peroxidation. This is in contrast with the findings of Spittle (2008).

Glutathione peroxidase reduces lipid hydroperoxides to their corresponding alcohols and free hydrogen peroxide to water respectively (Arthur, 2000). The major biochemical role of this enzyme is to protect the organism from oxidative damage. Our study showed that reproductive dysfunction is associated with reduced glutathione peroxidase activity. This observation is in contrast with the findings of Yamaguti *et al*. (2013) who observed an increase in serum glutathione peroxidase activities in rats administered continuously with NaF at varying concentrations. This observation however is in consonance with the studies of Shah and Khan, (2017) who observed a depletion in the glutathione peroxidase activity of rats induced with a well-established hepatotoxin. Treatment of reproductive dysfunction in male and female rats was not seen to reverse the alteration in glutathione peroxidase activity. This could be a result of a defective or impaired glutathione peroxidase activity as a result of a repressed expression or total inactivation of the enzyme since NaF is a well-established inhibitor of protein synthesis. This is in agreement with the findings of Agalakova and Gusev, (2012) who reported that fluoride changes the expression profile of apoptosis-related genes and causes endoplasmic reticulum stress leading to inhibition of protein synthesis. Pre-treatment with *T. indica* was however observed in our study to increase serum glutathione activity. This could be an indication of strong antioxidant property or enzyme inducing activity of the extract used in this study. This result can be compared to the findings of Shah and Khan, (2017) who reported an increased activity level of antioxidant enzymes and an ameliorated CCl_4_ after a preventive treatment with silymarin in combination with CCl_4_.

Glutathione is an important non-enzymatic antioxidant for neutralizing the toxic effects of some xenobiotics and in mopping up free radicals (Masella *et al*., 2005). At normal physiological state, the serum and cellular GSH concentration is abundant as it helps protect the cell from lipid peroxidation, serves as a regulator and activator of several transcription factors among its various roles (Masella *et al*., 2005). Since NaF compromises the serum activity of this soluble antioxidant as observed in our study and by Yamaguti *et al*. (2013), the result of our study in the treatment group suggests that *T. indica* may have inducing role in the production of GSH to counter the effect of free radicals produced during NaF induced reproductive dysfunction as an increase was observed in the serum concentration of GSH in all treatment groups compared to the negative control; thus restoring the GSH activity.

Catalase is a vital enzyme that protects the cell from oxidative damage by reactive oxygen species (ROS) majorly by catalyzing the decomposition of hydrogen peroxide to water and oxygen (Chelikani *et al*., 2004). The result obtained in this study indicates that catalase activities increases with increased lipid peroxidation in the male and that it is associated with decrease lipid peroxidation in the female. Our result is in agreement with the findings of Morales-Gonzalez *et al*., (2010) who observed an increase in serum catalase levels in rats administered NaF at varying concentration due to increased oxidative stress caused by the NaF induced reproductive dysfunction. The serum catalase activity in the pre-treated groups was comparable with the normal control group and this could be an indication of a strong antioxidant activity of *T. indica*. Treatment with the extract after induction of the dysfunction had a slight reduction in the serum catalase activity compared with the negative control. This could be a result of an increased susceptibility to proteolysis by degradation by specific proteases; an effect of NaF administration observed by Agalakova and Gusev, (2012).

Superoxide dismutase (SOD) is a vital antioxidant defense in nearly all living cells exposed to oxygen by catalyzing the dismutation of superoxide anion (Miller *et al*., 1990). In the present study, SOD activities in the male rats compared favorably with that of the female group; the increase observed in both groups is in agreement with the observation of Morales-Gonzalez *et al*. (2010) and Yamaguti *et al*. (2013) who also observed an increase in serum levels of superoxide dismutase in rats administered with NaF at varying concentration. Administration of *T. indica* prior to induction of sexual dysfunction and after inducing reproductive dysfunction also raised SOD activity. This observation is in contrast with the decreased toxicity and increased activity level of SOD in the preventive treatment carried out by Shah and Khan (2017).

## CONCLUSION

The importance of herbal products in the treatment of infertility in developed countries as well as in developing countries is undeniable. The administration of the fruit pulp extract of *Tamarindus indica* was found to have both prophylactic and ameliorative effect on oxidative stress parameters in sodium fluoride induced reproductive dysfunction in Wistar rats as evidenced by the oxidative stress parameters result. The most remarkable effects were observed with the 400 mg/kg body weight preventive and treated groups when compared with the control groups. Thus, the administration of the fruit pulp extract of *Tamarindus indica* at a dose of 400 mg/kg body weight could be used as an alternative to conventional drugs and methods in the management of oxidative stress in infertility. In addition, clinical and representative studies should be encouraged in order to complete the evaluation of the potential effects of *Tamarindus indica* in the treatment of reproductive disorders.

